# Prediction of fMRI activity using vector autoregressive models: a comparison of sparse and low-rank approaches

**DOI:** 10.64898/2026.06.11.731556

**Authors:** Xinle Tian, Alex Gibberd, Sandipan Roy, Matt Nunes

## Abstract

Vector autoregressive (VAR) models have a history of being used to examine functional connectivity in the brain, as captured by functional MRI studies. Such models allow for an estimation of Granger-causal relationships between regions of interest across the brain. Unfortunately, since the number of parameters in the VAR model scales as the square of the number of regions, and this is typically large compared to the number of temporal observations, these parameter estimates will exhibit high variance. To address this challenge, we introduce a low-rank pre-smoothing method that applies a low-rank approximation to the observations before fitting a VAR model. We estimate these models using individual subject data from both task-based and resting-state conditions, tuning hyperparameters at the population level. Our low-rank approach is directly compared against sparse and unconstrained estimation methods. Evaluations of predictive performance and model structure reveal that our pre-smoothing technique enables robust individual-level parameter estimation and significantly reduces prediction error, a finding further validated by synthetic experiments where the ground-truth parameters are known.

## 1 Introduction

Understanding dynamic dependencies in high-dimensional time series is a central challenge in modern statistical and neuroimaging applications [1, 2]. In particular, vector autoregressive (VAR) models provide a natural framework for capturing temporal interactions among multivariate signals [3–6], such as brain activity measured with functional magnetic resonance imaging (fMRI) techniques [7, 8]. However, classical VAR estimation becomes unstable when the number of variables measured is large relative to the sample size, a common situation in neuroscience and other domains involving high-dimensional data. This motivates the development of methods that improve estimation accuracy while preserving interpretable temporal relationships.

A natural extension is the use of sparse VAR (sVAR) models with ridge or lasso penalties [9], which regularise estimation and eliminate unnecessary connectivity among brain regions of interest (ROIs). While such approaches provide stability, they are inherently limited in their ability to capture shared latent dynamics across subjects, leading to a loss of interpretability at the population level. An alternative perspective emphasises low-rank structure in the VAR coefficient matrix, reflecting the presence of a smaller number of latent dynamical processes driving high-dimensional observations. Low-rank representations can capture shared dynamics and reduce variance, but may introduce bias by enforcing global structure that conflicts with localised interactions.

In this work, we focus on contrasting low-rank and sparse strategies for VAR estimation. Rather than imposing structural constraints or penalties directly on the coefficient matrix, we consider a procedure that modifies the input to the estimation objective. The central idea is to pre-smooth the response matrix through a low-rank projection, which captures latent temporal structures shared across variables. Although this procedure does not rely on explicit prior assumptions about the true VAR parameters, it typically yields coefficient estimates with an implicit low-rank structure. This setting allows us to examine low-rank and sparse modelling approaches side by side, and to study their respective implications for bias, variance, and interpretability in high-dimensional time series models.

From the perspective of fMRI analysis, our interest in deploying a low-rank procedure is motivated by the many existing applications of low-rank approximation applied to neurological data. For example, Independent Component Analysis (ICA) is often used as a pre-processing step to identify latent factors (or signals) driving the high-dimensional fMRI observations [10], and of course Principal Component Analysis (PCA) is ubiquitous in both vizualisation and as part of analysis pipelines. Let us denote the recorded fMRI activity at time *t* as a vector *y*_*t*_ *∈* ℝ^*p*^, where *p* is either the number of ROI or voxels. With ICA, the aim is to approximate the data *Y* ^*T*^ = (*y*_1_, ..., *y*_*n*_) according to the decomposition *Y* ^*T*^ *≈* Λ*F*, where *F ∈* ℝ^*k×n*^ represents a set of *k* (typically less than *p*) latent factors (sources), and the loading matrix Λ *∈* ℝ^*p×k*^ determines how each observation is described by these factors. Depending on the constraints imposed on the loadings, one may or may not interpret these factors. For example, if one imposes both orthonormality constraints and non-negativity conditions on the estimated 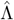, these loadings are identifiable and can also be interpreted as ‘networks’ (more accurately communities) where nodes *i ∈ {*1, ..., *p}* are connected if Λ_*il*_ *≠* 0, and there are *l* = 1, ..., *k* such communities [11, 12]. For example, a comparison of such low-rank approximation methods (applied to investigate brain-age prediction) can be found in [13].

We note that whilst low-rank approximations of *Y* have been extensively considered for fMRI applications, these typically do not consider direct analysis of the autocorrelation structure that may be present, i.e., the focus is on spatial/regional correlations, rather than the temporal dimension. Lately, there has been an increasing interest in fMRI activity forecasting, for example see the work of [14, 15], either modelling activity at the voxel, or ROI level. Whilst we acknowledge recent contributions in neural-network based forecasting, e.g. [15, 16], our focus here is on *efficient and interpretable* latent representation of series, which is difficult to achieve with highly non-linear methods. In fMRI forecasting tasks, modelling the latent dynamics in the series can be challenging due to the high levels of observational noise which masks the underlying structure. One way to cope with data with high noise settings is to use so-called pre-smoothing, in which response data is smoothed in some way prior to model-fitting [17, 18]. The motivation behind the first step in this approach is reduce the variance of the noise which effectively reduces parameter uncertainty. Pre-smoothing has been shown to benefit estimation and prediction in regression, see e.g. [19–21] and facilitate improved discovery of latent data structure [22–24]. The technique of pre-smoothing has recently been extended to the multi-response regression setting [25].

In this work we examine how the simple low-rank pre-smoothing approach of [25] can be combined within a VAR modelling framework to enable forecasting of fMRI activity. Our contributions are first to show the efficacy of our LRPS approach through synthetic simulations (with varying ground-truth structure, that is known), and then deploy the method on a variety of fMRI datasets to illustrate the prediction performance and empirical behaviour of the resulting estimated models. In particular, we contrast our approach with sparse VAR methods [9, 26, 27]. We also study performance across a cohort of individuals, to assess not just average performance, but the quantitative and qualitative similarity of estimates across subjects.

We structure the article as follows. Section 2 presents the methodology, including background on VAR models, sparse regularisation approaches, our proposed LRPS method and related works. Section 3 discusses the results from synthetic experiments comparing our proposed method with conventional methods. Section 4 presents results on predictive performance and the estimated model structure obtained using real fMRI datasets. A detailed description of the datasets used in this study is provided in Subsection 4.1. Finally, Section 5 offers a discussion of the findings, and directions for future work.

## 2 Background and methods

In this work, we adopt the common approach of representing activity in the brain using a VAR model for each individual within a population. That is, for each individual *l* = 1, ..., *m*, we assume that the data is generated according to the VAR(*q*) model:

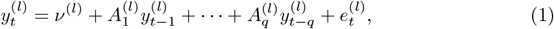

where *q* represents the order of the model, i.e. the number of time-lags incorporated into the model. Let *p* be the number of ROI in the dataset, we have 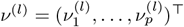 be a *p* vector of intercept terms (representing average activity) and the matrices 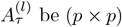 be (*p × p*) coefficient matrices that encode how regions interact at lag *τ* = 1, 2, ..., *q*. Finally, 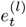 represents a noise vector (typically assumed to be independent and identically distributed across time and individuals) with mean zero and nonsingular covariance matrix 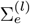.

In general, the order *q* of VAR models is often estimated via information criteria, e.g. via the Akaike IC (AIC) or Bayesian IC (BIC) [28, 29]. These criteria aim to balance the goodness of model fit (to data) and model complexity by penalising the log-likelihood in relation to the degrees of freedom in the model (number of parameters to be estimated). However, in neurological applications, according to previous studies, a first order model (*q* = 1) is often suggested to be the most relevant option for the VAR model with application to fMRI datasets [5, 30, 31]. Whilst the order of the VAR model used can significantly impact performance (either in an exploratory or predictive sense), deciding on this order is not the main objective of this work. Rather, we take the VAR(1) model and explore a simple method aimed at reducing the variance of the parameter estimates. Our decision to focus on the VAR(1) here is due to the inherent high-dimensionality involved with fMRI data, where one typically has *n ∈* [100, 200] time-points, and *p ∈* [10, 200] regions of interest. In this setting, without additional restrictions, the number of free parameters in the VAR(1) will be of order *p*^2^, and is often larger than the sample size *n*.

### 2.1 Least squares estimation

We will be primarily interested in the structure of the matrices *𝒜* := *{A*^(*l*)^*}*_*l∈𝒫*_, where *𝒫* represents a population of subjects. In our case, these subjects will be undertaking similar tasks/activities across the period of time for which the fMRI data is gathered. In what follows, we want to try to estimate the matrices at an individual level, i.e. not borrowing information across subjects; however, ideally, we will also be able to make some inference on the regions (and interactions) which are important at the population level. Before detailing our approach to this, we recap how one may classically estimate a VAR model using the method of ordinary least squares (OLS).

For notational convenience, we will here focus on the VAR for a particular individual and temporarily (without loss of generality) discard the ^(*l*)^ index. Suppose that we have data across *p* ROIs, and *n* time-points for this individual, then we can consider the matrices

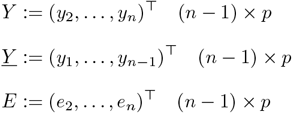

and for *t* = 2, ..., *n*, the VAR(1) model can be written in a compact form as

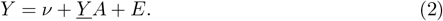

To simplify our discussion, and without loss of generality (as we can simply subtract the sample mean from each ROI) from herein, we assume *v* = 0.

In general, maximum likelihood estimation can be used to obtain the parameter estimates, based on an assumed distribution for the noise process *{e*_*t*_*}*. However, for simplicity, we consider the ordinary least squares (OLS) estimator defined as

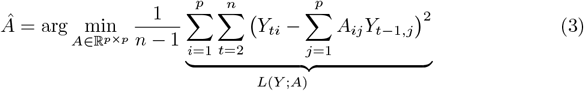

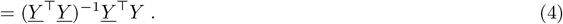

In the case where the noise is assumed Gaussian, with Σ_*e*_ = *I*_*p*_ then the OLS estimator is equivalent to the maximum-likelihood estimator (MLE).

Substituting (4) into (2) we note that the error incurred by the OLS estimator is given via the decomposition

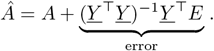

If one assumes *e*_*t*_ is independent of *e*_*s*_ for *s* ≠ *t* (and all fourth moments of the error exist and are bounded), and the true VAR process is stationary, i.e. the maximum eigenvalue of the VAR matrix is bounded *γ*_max_(*A*) *<* 1, then one can analyse this error [32, 33] to find

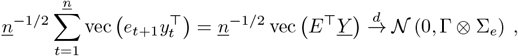

where 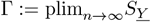 where 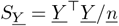 and such that plim denotes as convergence in probability with 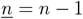, and the convergence in distribution occurs asymptotically as *n → ∞* [34, 35]. *As a consequence, asymptotically the OLS estimator (4) is unbiased and consistent, i*.*e*. 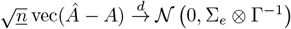

### 2.2 Regularisation and model selection

*Whilst the above results hold when n → ∞*, in reality, we have a finite *n*, with *p* of a similar order. When *p > n* we will not be able to directly invert the empirical covariance estimator 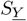, and thus one needs to provide an alternative estimator to the OLS. To address this challenge, two broad strategies have been proposed: restricting the parameter space, or transforming the data prior to analysis. The first approach includes regularisation techniques such as LASSO and ridge regression, which introduce penalties to stabilise estimation and reduce variance [9, 36]. The second approach involves dimension reduction techniques that operate on the data itself, such as using principal component analysis to extract lower-dimensional signals, e.g. performing a form of factor analysis [37, 38]. Both methods aim to impose structure or smoothness on the model, effectively balancing the trade-off between bias and variance and enhancing the predictive performance and interpretability of high-dimensional VAR models.

Regularisation/penalised estimators typically take the following form:

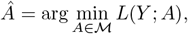

for some restricted subspace *ℳ ⊂* ℝ^*p×p*^, where typically *Â*_*OLS*_ ∉ ℳ. An alternative view of these estimators is from the penalisation (or so-called Lagrangian) perspective:

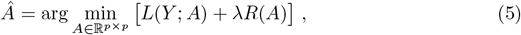

where *λ ≥* 0 penalises the loss function *L*(*Y* ; *A*) by scaling a penalty function *R*(*A*). The form of the function *R*(*A*) is chosen to encourage *Â* to possess some desirable properties, e.g., we can use *R*(*A*) = *∥A∥*_1_ := Σ_*ij*_ |*A*_*ij*_| to encourage sparsity within 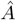 [36, 39, 40], or 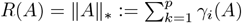 (the sum of the eigenvalues) to encourage a low-rank *Â* [41]. One can also note that classical information criteria such as AIC/BIC are also written in the form of (5), however, these involve the addition of non-convex counting penalties, which typically make solving the minimisation problem computationally intractable.

As an alternative to our proposal, we will compare our new approach with the existing and popular sparse-VAR (sVAR) model, given by [9]

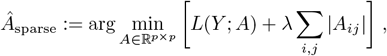

where *λ* can be selected based on either information criteria, or cross-validation based approaches.

### 2.3 Low-rank pre-smoothing for VAR estimation

In this section we introduce an application of the Low-Rank-Pre-Smoothing (LRPS) estimator developed in [25] to estimation of parameters in the VAR model. Our proposal is straightforward: rather than searching for the best low-rank representation (e.g. via an M-estimator of the form Eq (5)) we simply project the data into a certain subspace and then perform OLS estimation in that space.

Specifically, let *Y* = *UDV* ^*T*^ be the singular value decomposition (SVD) of *Y*, then we approximate the data by retaining the top *k* components:

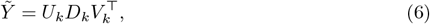

where *D*_*k*_ is a *k × k* diagonal matrix with the first largest *k* diagonal entries of *D. U*_*k*_ and *V*_*k*_ are *n × k* and *p × k* matrices by keeping the *k* corresponding columns in *U* and *V* . In choosing the approximating rank 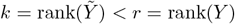, we need to consider a trade-off in the reduction of variance (due to reducing the dimensionality) and an increase in bias (as 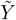 is an approximation to the original data). Although a small *k* is usually required in the graphical representation of the whole dataset [42], there are circumstances in which more factors are needed.

To model temporal dependence, we now consider how the lagged observations 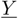 drive this approximate data matrix 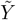 . This motivates us to consider the regression

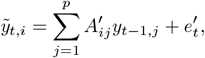

where we note *A*^*′*^ and *e*^*′*^ are distinct from *A* and *e* in the original VAR model (2), as we are now working with the approximation. Note that (6) can also be written as 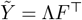 where 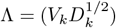 and 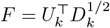. In matrix notation, we have

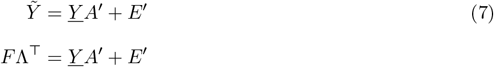

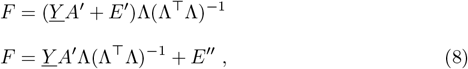

and thus we can equivalently study the regression with *p* outcomes, or in the latent (factor) space with *k* outcomes, where we have *A*^“^ = *A*^′^Λ(Λ^*T*^Λ)^*−*1^ *∈* ℝ^*p×k*^ . We further note, that with our choice of using the SVD for pre-smoothing, that Λ^*T*^Λ = *D*_*k*_ due to the columns of *V*_*k*_ being orthonormal. In terms of estimation, we propose to estimate *A*^*′*^ using OLS (the same as in [25]) resulting in the LRPS estimator

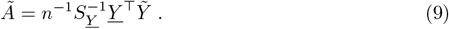

We propose to use this estimator Ã as a plug-in estimator for the original VAR model (2), and consider its ability to recover the true *A* (not *A*^*′*^), and the predictive performance of the resulting VAR model.

### 2.4 Multi-subject cross-validation procedure

As one may expect (and illustrated in our experiments later), the choice of *k* is critical in achieving good finite-sample performance with LRPS. To make this choice in practice, here we develop a cross-validation (CV) framework that operates across multiple subjects/trials. In our applications, we also deploy this method to choose the regularisation parameter *λ* for the sparse-VAR approach.

Given that the VAR model is predictive in nature, it is natural to adopt a framework based on evaluating predictive performance. In this case, we consider a *rolling forecasting origin*, where the training set (for each subject) expands sequentially. The process begins with an initial training set of a specified size, making predictions for the next time point. Then, the training set expands by including the next observation, and a new forecast is made for the subsequent point. This rolling process continues, ensuring that only past data is used for prediction, mimicking real-world forecasting constraints. Forecast accuracy is evaluated by averaging errors across all test sets, providing a reliable measure of model performance over time. We define the cross-validation mean squared prediction error (cvMSPE) for hyperparameter *k* as:

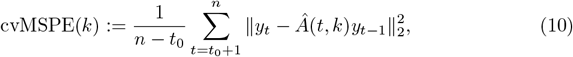

where *n − t*_0_ is the total number of splits, i.e., the number of rolling window iterations used for cross-validation. For each *t, k* we obtain a potentially different estimate *Â*(*t, k*), where the goal is to find the best hyperparameter 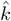 that minimises the cvMSPE

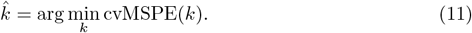

Note, we effectively replace *k* with *λ* in this framework to tune the level of shrinkage in the sparse-VAR estimator.

Because fMRI datasets often include recordings from multiple subjects, it is important to design a CV strategy that accounts for inter-subject variability. To address this, we adopt a multi-trial CV framework, where each subject is treated as an independent trial. Within each subject, we perform rolling-origin time series CV to evaluate model performance across different parameter choices, e.g. the rank *k*. The optimal value is selected for each subject based on minimising the MSPE, and a consensus *k* is determined by aggregating the subject-level selections, such as by taking the most frequently chosen rank across subjects (see Figure 1 for a diagrammatic representation of this framework). This ensures that the final hyperparameter generalises well across the population, not just within individual subjects. In general, choosing a single hyperparameter across a population can help us when interpreting the results from model fitting, as each subject will have had its model restricted in a comparable way.

**Fig 1.**
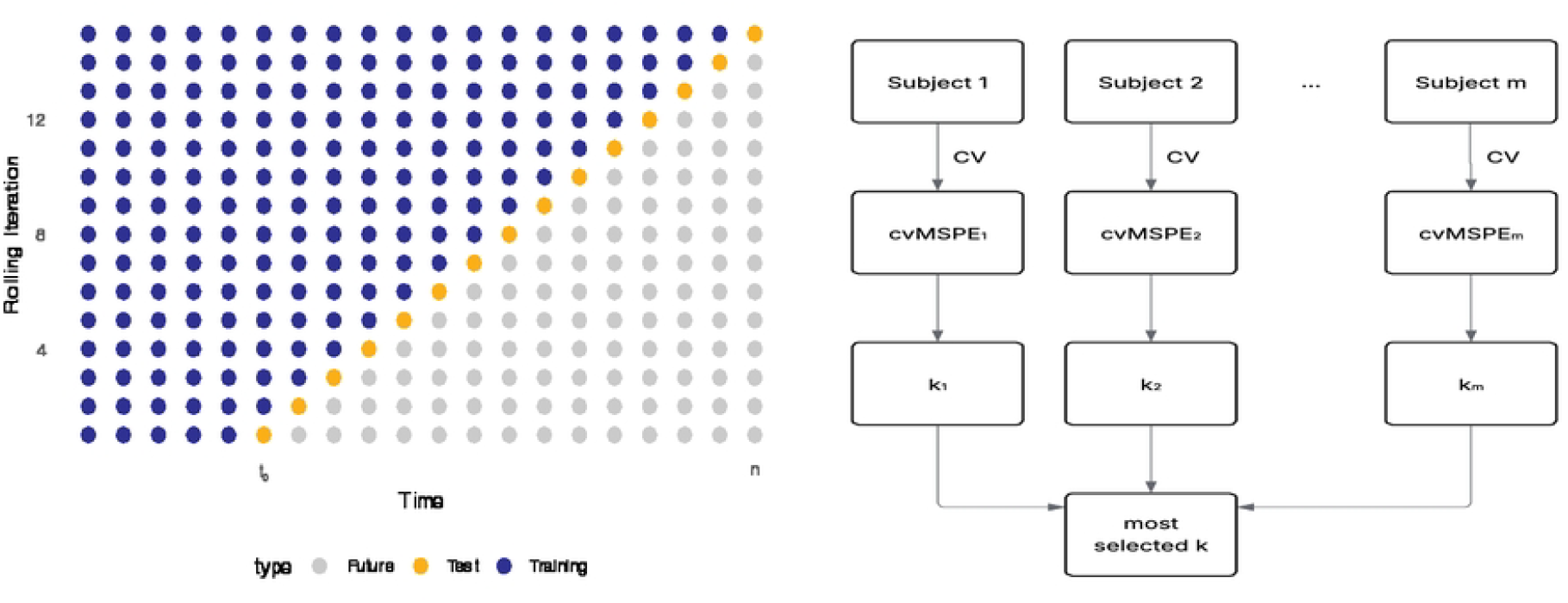
Visualisation of multi-trial cross-validation. Left: rolling-origin time series cross-validation performed on a single subject, where the model is iteratively trained and evaluated on temporally advancing windows to estimate MSPE. Right: multi-trial CV framework applied across multiple subjects, where each subject undergoes the same rolling CV process to compute MSPE across different *k* and the final parameter is based on aggregating the optimal *k* values across subjects.

### 2.5 Granger causality

One important feature of the VAR model is its capacity to facilitate Granger causality analysis [43], which aims to determine whether the past values of a given node *i* contain predictive information about the future values of another node *j* [44]. In the context of networked time series such as brain activity, Granger causality provides a valuable tool for uncovering directional relationships between regions [45]. When we are in a high-dimensional setting, i.e. *p ≈ n*, the sparse-VAR method of (2.2) is useful as it is computationally tractable, has the ability to strike a bias-variance tradeoff via the adjustment of *λ*, moreover, it explicitly produces zero estimates in the *Â* matrix. One can interpret the estimated non-zero coefficients as encoding a form of Granger causality “network”, thus the sparse regularisation approach naturally helps us uncover the structure in this network.

Given the low-rank constraints effectively imposed by LRPS, extraction of sparse Granger casual networks is more involved, since we would typically expect the estimates *Ã* to be dense (i.e, not have many zero values). However, we can see by comparing Eqs. (7) and (8), we could alternatively have estimated *A*^*′′*^ in the dimension-reduced space according to 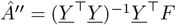 and then invert the relationship *Â*^*′′*^ = *Â*^*′*^Λ(Λ^*T*^Λ)^*−*1^ to obtain *Â*^*′*^ = *Â*^*′′*^(Λ^*T*^Λ)Λ^*T*^(ΛΛ^*T*^)^*†*^ = *Â*^*′′*^Λ^*T*^ where *†* denotes the Moore-Penrose inverse. With this in mind, a form of Granger causality interpretation may be considered more appropriately at the factor level, where we see which of the lagged observations drives which of the *l* = 1, ..., *k* factors by looking for large values in the rows of *Â*^*′′*^. To be clear, we cannot prescribe a direct Granger causal relationship based on *Â*^*′′*^, as the factors *F* are linear combinations of all ROI’s, however, the structure in *Â*^*′′*^ would allow us to see which regions in the brain are *driving* these factors, from the previous time-step.

### 2.6 Related work

In the context of fMRI data analysis, the LRPS approximation 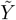 can be related to the idea of global signal regression (GSR) [46, 47] where regional activity is regressed onto the mean activity of the brain (at each point in time). However, in our case, we have a dynamic (lagged) regression, and we would be investigating which regions drive the global signal. For example, if we considered that *k* = 1 and assumed that 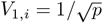, then the corresponding factor *F*_*t*,1_ could be considered as the global (mean) signal. The regression (8) then looks at how each lagged region drives this global mean signal. Again, for *k* = 1, we would have *A*^*′′*^ *∈* ℝ^1*×p*^. If the value of 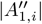 was large, then it could indicate that the *i*th region had a significant role in driving the global signal at the next time-point. Importantly, in our case, the columns of *V*_*k*_, *v*_*l*_ for *l* = 1, ..., *k*, are not explicitly chosen to represent the global signal, and instead are data-adaptive, i.e., *V*_*k*_ is specified via the SVD. This aligns somewhat with the work of [48] who propose a low-rank approximation approach to correlation maps (based on seed locations), and show when the data is clustered around their first principal component, the correlation maps become dominated by the GSR signal. In our case, if a global signal is present in the fMRI data, one may expect *v*_1_ to be fairly flat; however, we do not force the presence of this structure. We also do not base our analysis around a particular seed region; the importance of a specific region in the estimation is given via the loss function, and is ultimately based on the variation of each ROI’s time-course. Our approach is also somewhat related to methods that perform dimensionality reduction and then assess Granger causality in that reduced space, such as dynamic factor models [37], or Granger PCA [49] approaches. These frameworks operate under the assumption that a small number of latent factors or principal components drive the dynamics of the observed variables, and they seek to model the temporal dependencies among these hidden components, e.g. a VAR model may be specified in the latent space. The main difference between our approach and these is that we regress the projected data 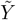 onto the original (not approximated) lagged variables with the aim of assessing which regions of the brain are driving the factors at the next time-point.

## 3 Synthetic experiments

In this section, we assess the performance of the proposed LRPS method, VAR and sVAR models under a range of controlled synthetic experiments. We focus on scenarios where the structure of the coefficient matrix *A* is explicitly specified and known, allowing us to systematically explore how each method performs when *A* exhibits varying structures. We also consider the effectiveness of our parameter tuning method as detailed in the previous section, and in preparation for real-data analysis.

### 3.1 Simulation settings

To keep the analysis straightforward, we focus on the case of a first-order lag in our simulations. The data are simulated according to (2), where the errors are assumed to be independently generated according to a standard normal distribution. We set *p* = 30, *n* = 100 and we vary the specification of the true coefficient matrix *A*. The choices here cover a range of scenarios that are of interest in practice and help further elucidate the behaviour of LRPS and VAR when the spectrum (eigenstructure) of the *A* varies. Note that simulations in this section are performed in the *R* statistical computing environment [50]. More specifically, data are generated by *tsDyn* package [51] and VAR fitting is through *vars* package [52] (using OLS). The various conditions for *A* are given below:

1. *Flat eigenvalues:* The matrix *A ∈* ℝ^*p×p*^ is constructed as a matrix where the diagonal elements are set to *ρ* and the off-diagonal elements are set to either 0.2 or 0.01 (referred to as Experiment 1A, and 1B respectively) to examine how different magnitudes of eigenvalues impact the performance of models:

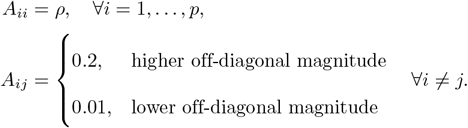
2. *Decaying eigenvalues:* In this case, we investigate the setting where the dependence decays from the diagonal (geometrically) according to:

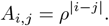

and speed of decay in the eigenvalues is modulated via *ρ*. To ensure stationarity, if *σ*_max_ *≥* 1, for an experimental configuration, the VAR matrix is rescaled to maintain stationarity, i.e. *A → A/*(*γ*_max_(*A*) + 0.1). We see that when *ρ ≈* 1 that the *A* matrix is very ill-conditioned, i.e. *γ*_max_(*A*)*/γ*_min_(*A*) will be very large. In this setting, the matrix possesses a lower effective rank [53], and may be approximated as a low-rank matrix.

Figure 2 shows how the spectral properties of the matrix *A* change with different values of *ρ* (0.1, 0.5, and 0.9) in each of the simulation settings. In the cases of the flat structure (both red and blue lines), we see that the first eigenvalue is significantly larger than the others, and the spectral gap (jump between the first and second eigenvalues) is moderated by both the scale of the off-diagonal elements, i.e. contrast high and low off-diagonal settings, and the on-diagonal scale of *ρ*. A larger *ρ* results in a smaller gap. In the case of a decaying structure (green line), we see that a larger *ρ* leads to faster eigenvalue decay, indicating that the structure of *A* becomes more concentrated in a few leading components. For lower *ρ*, the decay is more gradual, suggesting that a higher-rank approximation may be needed.

**Fig 2.**
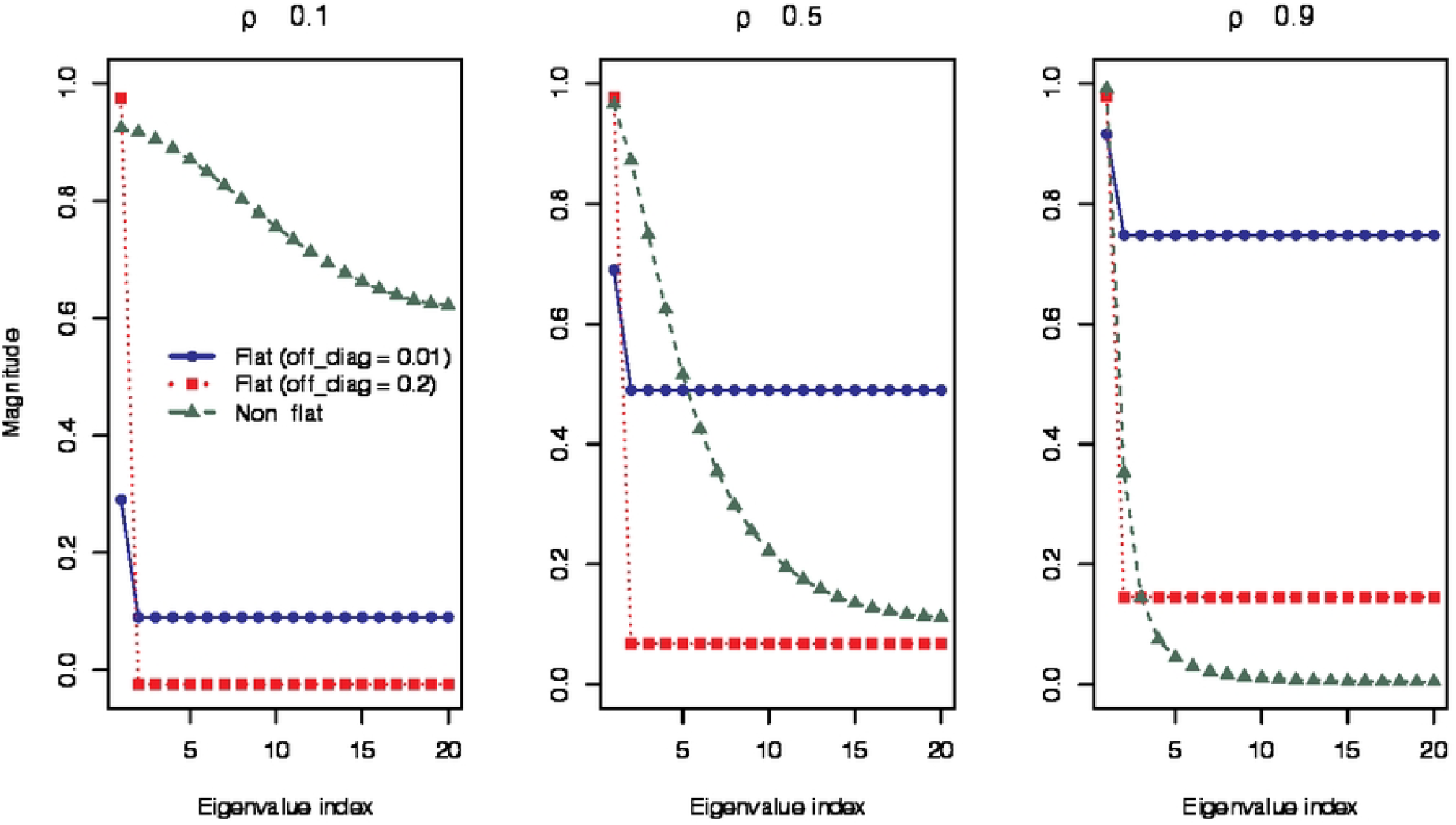
Eigenvalue structure for selected cases. Red squared line: Experiment 1A - flat structure with higher off-diagonal magnitude. Blue circle line: Experiment 1B - flat structure with smaller off-diagonal magnitude. Green triangle line: Experiment 2 - geometrically decaying structure.

### 3.2 Results – estimation error

As we have access to the ground-truth *A*, we can assess the estimation performance directly, for which we use the mean squared error (MSE) defined as

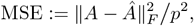

where *∥A∥*_*F*_ = Σ_*ij*_ |*A*_*ij*_|^2^ denotes the Frobenius norm and *Â* is the estimated coefficient matrix (for each method under consideration). Each scenario is replicated 100 times, and the average MSE is computed for the final performance evaluation.

Figure 3 presents the MSE results as a function of the relevant tuning parameter, under varying values of *ρ* for the experimental scenarios. In all cases, the VAR model maintains a constant MSE value, which aligns with the fact that there is no rank selection or tuning parameters involved. The plots demonstrate the behaviour for both the LRPS estimator and SVAR, with their respective tuning parameters *k* and *λ* given on the horizontal axes.

**Fig 3.**
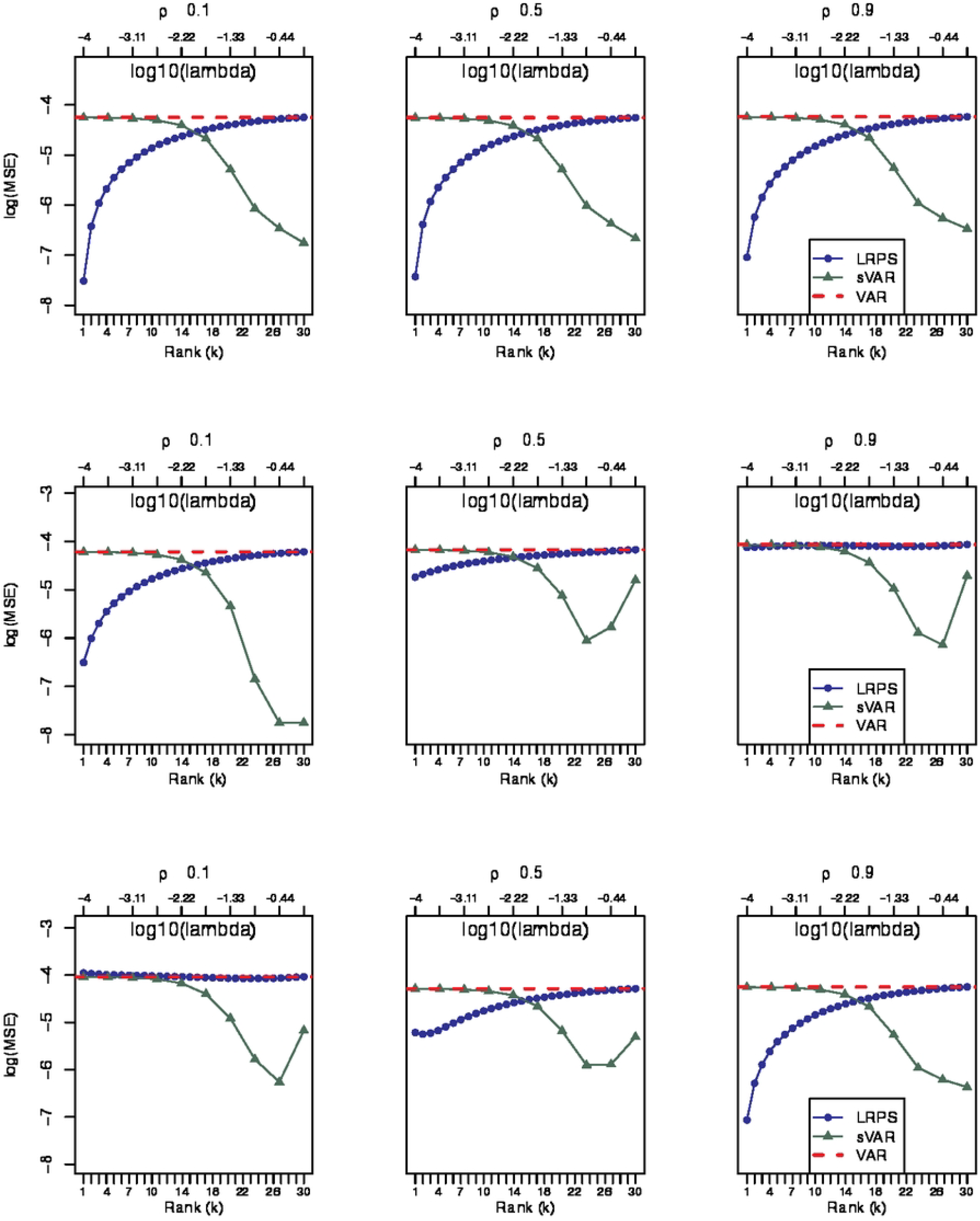
Log-MSE results from synthetic experiments with LRPS (blue), VAR (red), and sVAR (green) for *ρ* = 0.1, 0.5, 0.9. Top: Experiment 1A; flat structure with higher off-diagonal magnitude: *A*_*ij*_ = 0.2, *∀i* ≠ *j*. Middle: Experiment 1B; flat structure with smaller off-diagonal magnitude: *A*_*ij*_ = 0.01, *∀*i ≠ *j*. Bottom: Experiment 2; geometrically decaying structure on the true *A*.

In contrast to the OLS estimator the LRPS method exhibits a rank-dependent behaviour, as the rank of choice *k* increases, the MSE of LRPS gradually approaches the VAR model. At lower ranks, these methods impose stronger dimensionality constraints, which may result in lower MSE due to improved bias-variance trade-offs, especially when the true coefficient matrix has a low-rank structure. However, as the rank increases towards full rank, the constraints are relaxed, leading to a convergence towards the performance of the standard VAR model. We see that when the spectral gap is large, the difference between the low and full-rank (VAR) estimates is much greater, e.g., we can relate the rows of results in Figure 3 with the jump in eigenvalues in Figure 2.

The sparse VAR also demonstrates competitive performance in these experiments, especially when the off-diagonal elements are well approximated as zero, namely experiments 1B, and 2, the latter in the cases where *ρ* is small, so the geometric decay is quickly towards zero. Ultimately the best method to use depends on the structure in the true *A* matrix, however, the experiments illustrate how both LRPS and sVAR allow us to adaptively improve estimation of VAR models.

### 3.3 Results – prediction error and rank selection

Whilst low estimation error of parameters will in this case (as the model is well-specified) relate to good out-of-sample predictive performance, it is useful to assess the ability of our cross-validation procedure to appropriately select tuning parameters for the LRPS procedure. To this end, we consider a setup where *t*_0_ = *n −* 50, and thus 50 samples are used to gauge the predicition error associated with a grid of *k* values. After this procedure, 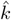 is selected according to Eq. (11) and a further 50 samples are used to evaluate prediction performance in an evaluation dataset (with the same generating process).

Figure 4 illustrates the distribution of selected rank across 50 simulation rolling windows, under conditions of Experiment 2 (Toeplitz form). For each time, the model selects a single *k* so the histogram shows how frequently each rank is chosen. The 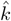 reported in the title denotes the multi-subject global optimal *k* identified using our multi-subject cross-validation procedure described in Section 2.4. In the density plots (Figure 4, second row), which illustrates prediction performance, this fixed global optimal *k* is used across all replicates. We see that depending on the true *ρ*, the approximating rank is automatically adjusted from being concentrated around 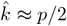, to being concentrated at a small number, e.g. 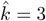 in the *ρ* = 0.5 case and 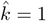 in the *ρ* = 0.9 case. This observation aligns with the eigenvalue spectrum in Figure 2: the more spiked the eigenvalue distribution, the more concentrated (and smaller) the optimal choice of 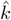 becomes. Similar results for Experiment 1A and 1B are presented in the Supplementary Information (5), the results show that when the off-diagonal magnitude is large, LRPS tends to select the rank *k* consistently small across simulations, where again in these scenarios, the true eigenvalue distribution is very spiked.

**Fig 4.**
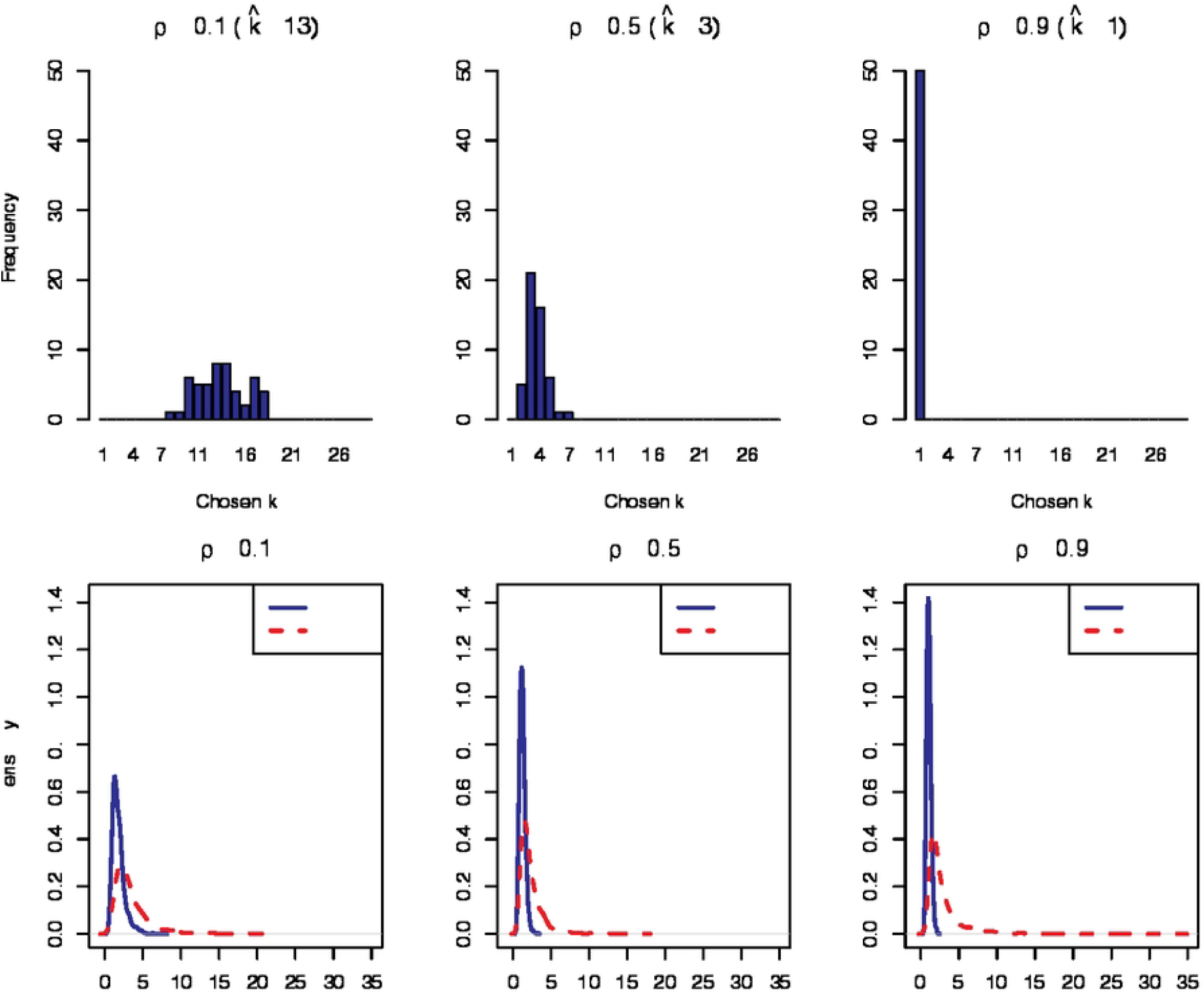
Results for rank selection and predictive performance under the conditions of Experiment 2. Top: Example of the hisogram for *k* over the 50 out-of-sample data-points; Bottom: Density estimate of the MSPE as estimated over the 100 replications.

Overall, we see that our cross-validation procedure does a good job at tuning the required approximating rank, with the predictive performance (using the cv-selected 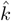) dominating the performance of the regular VAR model (as shown in the bottom panel of Figure 4). We note that the predictive performance is not only on average better, but also that there is less variability in this performance.

## 4 Dynamic modelling of fMRI ROI data

In this section, we compare the predictive performance and estimated patterns in the VAR matrix when applied to fMRI data, using the methods discussed in Section 2. In particular, we are interested in the LRPS method’s ability to reduce the prediction error associated with VAR models fit to data at an individual level. We contrast this with the performance achieved by a sparse-VAR model, where both the optimal *k*, and *λ* are chosen in a comparable manner. Finally, we examine structure (networks) that appear prominent after averaging the estimated VAR matrices across the population, and how the overall importance of the VAR *A* matrix (to describe brain activity) depends on the task individuals are asked to perform.

### 4.1 Experimental data

We use three distinct datasets in our study, all openly available from OpenNeuro, accessible at https://openneuro.org. We pre-process each dataset using the AAL atlas [54] (with *p* = 116 regions) and standardise the data for each subject. Following preprocessing, we exclude subjects whose time series data do not include a sufficient number of ROIs. The datasets and their sizes following pre-processing are summarised in Table 1. We give further details on the background for each dataset below.

**Table 1.** Summary of fMRI datasets analysed.

| Dataset | URL | Reference | #Sub | Size |
| --- | --- | --- | --- | --- |
| DMCC5B (Axcpt) | Link | [55] | 33 | $(610 \times 116)$ |
| DMCC5B (Cuedts) | Link | [55] | 35 | $(650 \times 116)$ |
| Rest | Link | [56] | 35 | $(160 \times 116)$ |

#### DMCC55B dataset

This dataset is a subset of the task fMRI runs collected as part of the Dual Mechanisms of Cognitive Control (DMCC) project. Detailed information about the dataset is provided in [57], while the theoretical framework and broader goals of the DMCC project are described in [58]. While the full dataset includes four different task paradigms, we focus on data from two tasks, namely the AX-CPT (Axcpt) and Cued Task-Switching (Cuedts) experiments to demonstrate the potential advantages of using LRPS in applications where the brain was subject to specific stimulii. Participants were asked to complete tasks, and respond based on pressing one of two buttons.

In the Axcpt task, participants respond to letter stimuli based on contextual cues. They pressed button 2 only if the current letter was X and the preceding letter was A; in all other letter conditions, they pressed button 1. Number stimuli serve as no-go trials, where participants are instructed to withhold a response. In the Cuedts task, each trial began with a cue instructing participants to either “attend number” or “attend letter”. The target stimulus consisted of a letter-digit pair presented side-by-side. If cued to “attend number”, participants performed an odd / even judgment (they pressed button 1 for even, button 2 for odd). If cued to “attend letter”, they performed a vowel / consonant judgment (they pressed button 1 for vowels, button 2 for consonants).

#### Resting-state fMRI dataset

To contrast with the above task-based activities, we also include a resting-state dataset in our analysis. This dataset consists of retrospective resting-state scans from patients diagnosed with brain tumors. We preprocess and standardise the data with the same method as previous datasets, selecting subjects with sufficient ROI coverage. Further details about the dataset can be found in [59] and [60].

### 4.2 Tuning parameter selection

To tune the sVAR and LRPS methods we use the multi-trial rolling forecasting cross-validation approach described in Section 2.4. For each task, we select the first 10 subjects and apply an initial training window. Due to differences in dataset sizes, we set the initial window to 300 time points for the DMCC55B dataset and to 150 time points for the resting-state dataset, the rest of the data for these subjects is used to assess the prediction error as per Eq 10. The resulting cross-validation performance can be seen in Figure 5. For LRPS, the candidate ranks are chosen from 1 to 10, as it will be computationally inefficient and unnecessary to evaluate the full range up to 115. Based on this procedure, the optimal rank selected for LRPS is 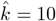 for the DMCC55B dataset in both the Axcpt and Cuedts tasks. In contrast, the optimal rank is 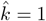 for the resting-state dataset. Note, although we present the cvMSPE for all methods, the tuning parameter 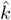 is based on a consensus (best *k*) estimate, and this may not directly align with the minimum cvMSPE averaged across all subjects, e.g. as reported in Figure 5 where we see the average error is minimized for 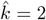. In practice, in this case, a choice of either 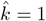 or 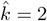 could be appropriate. For the sVAR model, we evaluate a sequence of regularisation parameters defined on a logarithmic scale, specifically with 20 values evenly spaced on a log scale from 10^*−*4^ to 1. The optimal value of *λ* selected for sVAR is 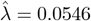 for both the DMCC55B dataset across both tasks and the resting-state dataset.

**Fig 5.**
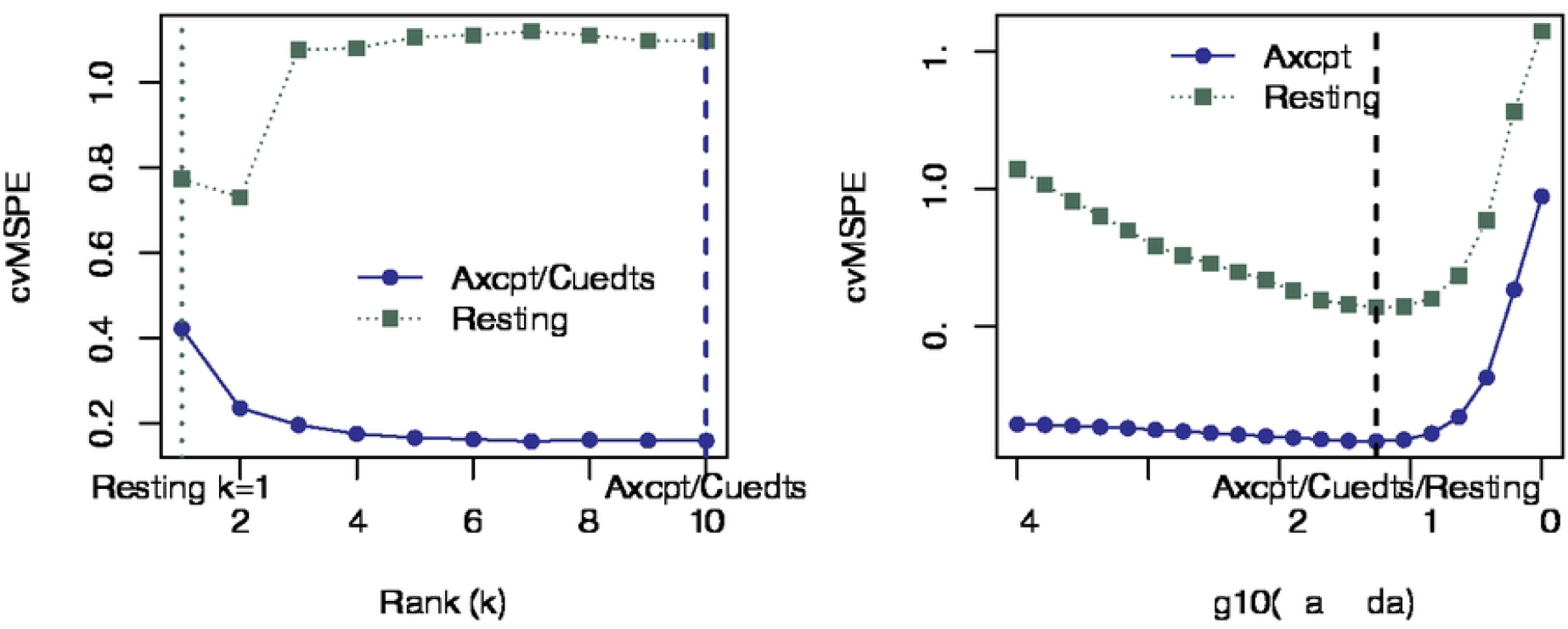
Cross-validation performance using a random selection of 10 subjects, with the left panel giving tuning results for *k* in the LRPS method, and the right panel giving those for *λ* with the sVAR approach.

We summarise the performance on the out-of-sample data (not used for this tuning) in the following sections. In particular, we are interested in the prediction error, and how the individual matrix *Â*^(*l*)^ vary across *𝒫* for the different methods.

### 4.3 Results and performance measures

For each of the datasets in question, we assess various aspects of the VAR model performance, for each of the estimators: LRPS, VAR, and sVAR. We are interested in both the predictive ability of the models, measured via mean-absolute prediction error (MAPE), and root mean squared prediction error (RMSPE) on a one-step ahead basis, where these predictions are gathered on out-of-sample data in that we consider the performance for each individual averaged across the last 10 time-points in the Axcpt and resting-state data, and 50 time-points in the Cuedts task-based experiment (which had the longest time-course). Note that similar to the simulation study in Section 3, this data is neither used to construct the estimates or select the hyperparameters (i.e. performance is taken over a distinct evaluation dataset); recall that for these data, the VAR models are estimated independently across the individuals involved in a dataset, however, tuning parameters in the LRPS and sVAR cases are set at a population level.

In addition to the predictive performance, we are also interested to see how the resulting VAR *Â*^(*l*)^ matrices are structured. To summarise causal (functional) relations at the population level, we simply average the estimated VAR matrices, and visualize

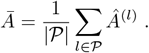

By considering the matrix *Ā*, we aim to average out individual differences in the predictive models, to gain some insight on interaction networks at the population level. Whilst this is a simplistic approach and can offer some insights, we must be careful that averaging over individuals can mask heterogeneous effects, for example, if individuals were clustered, it could be that when averaging across individuals it may appear as though there is no relationship between regions, i.e. *Ā*_*ij*_ ≈0, however, this may be the effect is averaged out across distinct sub-populations, that is we succumb to a form of Simpson’s paradox.

To gain some insight on the heterogeneity between estimates *Â*^(*l*)^ we consider a simple measure based on the principal angle between the individual estimates and the population average. For simplicity, we calculate this principal angle using the first eigenvector of *Â*^(*l*)^ for each subject and the first eigenvector of population average *Ā* [61]. A smaller principal angle indicates greater alignment between an individual subject’s estimate and the group-level estimate. If these angles are small, then one may have greater confidence in the interpretation due to averaging, that is via *Ā*. These angles also allow us some measure of how strong the individual differences within a task may be.

A summary of the performance of the three methods across the set of four experiments is given in Table 2.

**Table 2.** Summary of experimental results over fMRI datasets. The principal angle *π* represents the average of the angles presented in Figs. 6 and 7, whilst the average connection ave(*Â*) represents the average of the population VAR matrix, i.e. Σ_*ij*_ *Ā*_*ij*_ */p*^2^ for each method and dataset.

| Dataset | Method | RMSPE (SD) | MAE (SD) | ave($\pi$) | ave($\hat{A}$) |
| --- | --- | --- | --- | --- | --- |
| Axcpt | LRPS | 0.626 (0.254) | 0.480 (0.201) | **0.929** | 0.004 |
|  | sVAR | **0.466 (0.188)** | **0.341 (0.148)** | 1.036 | 0.005 |
|  | VAR | 6.979 (4.261) | 4.547 (2.648) | 1.365 | 0.005 |
| Cuedts | LRPS | 0.574 (0.191) | 0.436 (0.157) | **1.062** | 0.005 |
|  | sVAR | **0.449 (0.158)** | **0.330 (0.125)** | 1.241 | 0.006 |
|  | VAR | 6.688 (3.683) | 4.294 (2.175) | 1.363 | 0.006 |
| Resting | LRPS | 0.911 (0.316) | 0.738 (0.278) | 1.044 | 0.001 |
|  | sVAR | **0.633 (0.248)** | **0.488 (0.215)** | **1.018** | 0.001 |
|  | VAR | 18.984 (16.664) | 12.759 (11.208) | 1.362 | 0.001 |

Figures 6 and 7 give more detailed results on estimated structure and heterogeneity:

- The left column of the figures presents a summary of the estimated VAR matrices based on the remaining 23 subjects (not used for hyperparameter tuning), specifically the average estimated VAR matrix *Ā*. In each plot, we display the top 1% of connections across all ROIs to highlight the most prominent connectivity patterns. Although the absolute number of connections may differ across methods, the proportion remains consistent. It is also worth noting that this visualisation does not highlight self-connections, i.e. those resulting from on-diagonal structure in the estimated VAR matrices.
- The right plot shows group-level connectome maps, constructed by thresholding t-statistics at *α* = 0.01. Significant positive connections are shown in red and significant negative connections in blue, with colour intensity proportional to the magnitude of the t-statistic. Note that the t-statistic is simply calculated based on the entrywise mean divided by the standard deviation avg 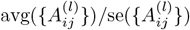 for each pair of ROIs *i, j*. For simplicity, we did not perform any adjustment for multiple testing when thresholding.
- On the bottom, we present a summary of the principal angles based on the first eigenvectors of *Â*^(*l*)^ and *Ā*. These give some indication of heterogeneity, and measure of how representative the population averages may be for the individual estimates.

**Fig 6.**
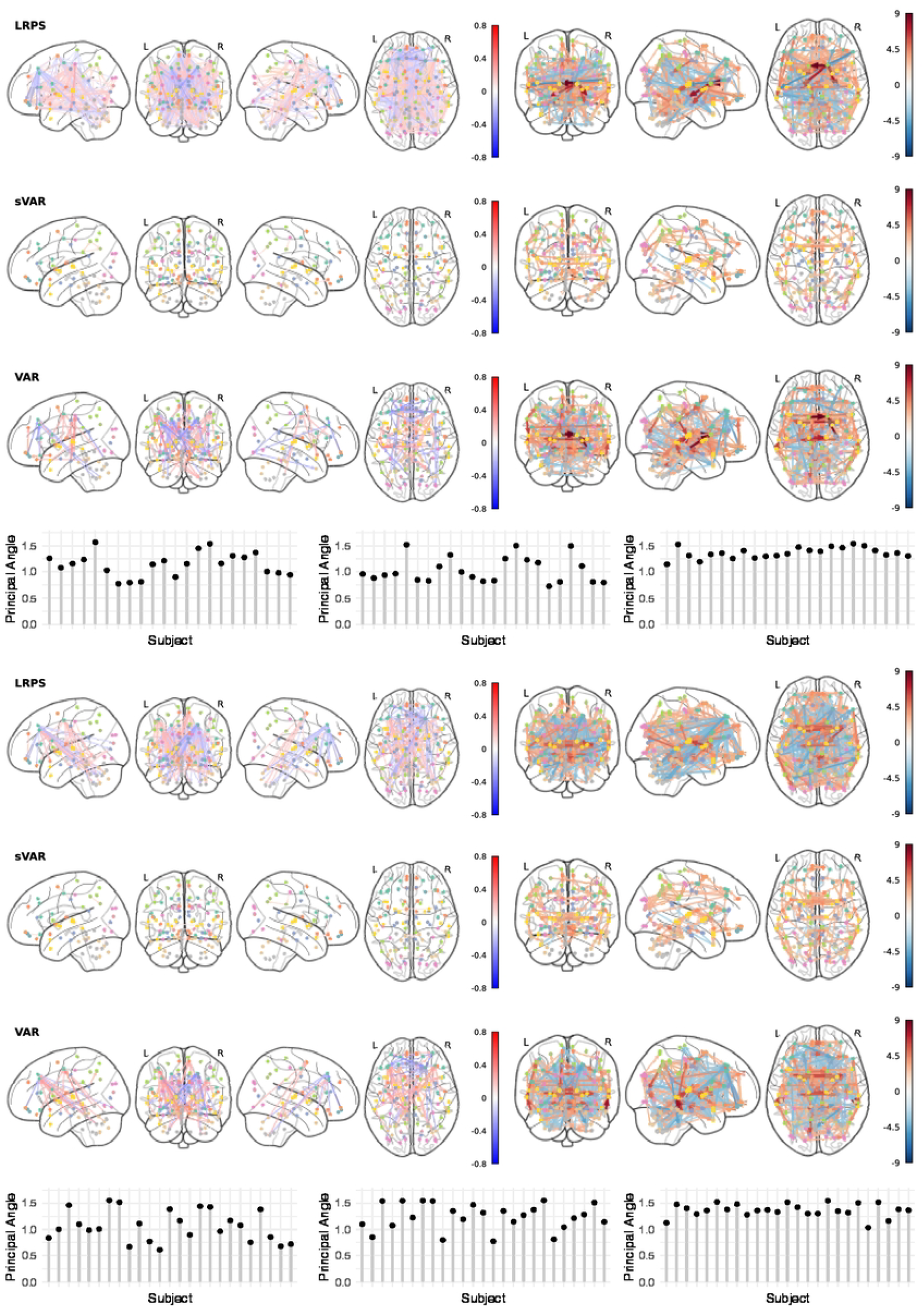
Results for the Axcpt task (top-panel) and Cuedts task (bottom-panel). Left: Visualization of average estimated VAR matrix. Right: Group-level connectome based on one-sample t-tests of estimated VAR matrices. Bottom: Principal angle for individuals in the evaluation set (LRPS, sVAR, VAR, left-right).

**Fig 7.**
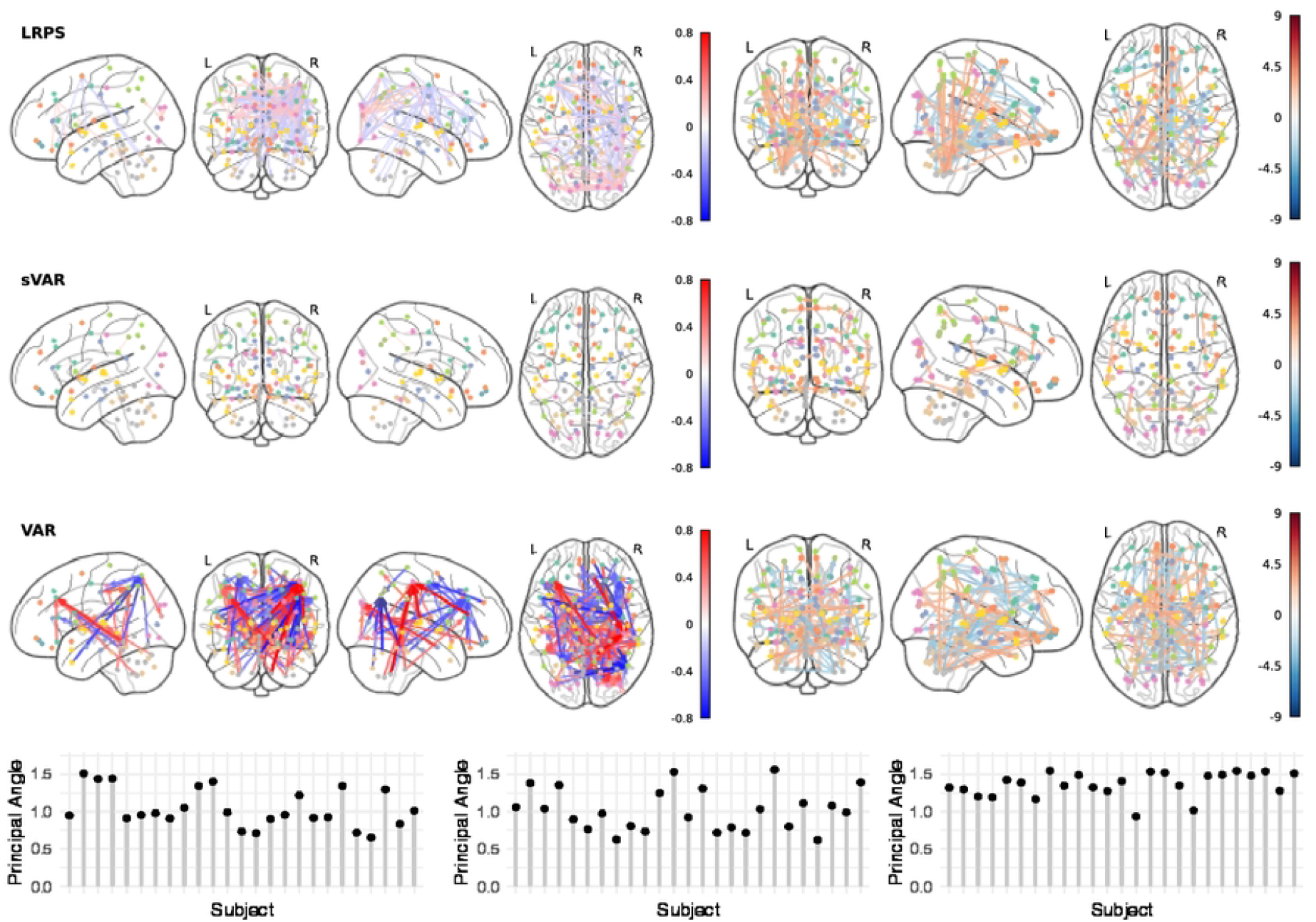
Results for the resting-state data. Left: Visualization of average estimated VAR matrix. Right: Group-level connectome based on one-sample t-tests of estimated VAR matrices. Bottom: Principal angle for individuals in the evaluation set (LRPS, sVAR, VAR, left-right).

### 4.4 Discussion

The empirical results in Table 2 demonstrate that the LRPS-VAR approach drastically improves over the regular VAR model in terms of the predictive performance. In the task-based experiments, the error is reduced relative to the VAR model by roughly a factor of 10, whereas in the resting-state application the reduction in the predictive error is even more significant (from an RMSE of 18.984 for VAR to 0.911 for LRPS). This improvement in predictive performance highlights the importance of striking a reasonable trade-off when constraining the flexibility of the model, and we see that LRPS provides a valid mechanism to do so. However, whilst LRPS does improve predictions relative to VAR, we see that the sparse-VAR approach (sVAR) improves even further. Indeed, sVAR wins in all scenarios considered here in terms of MAE and RMSE performance. The gains in performance are significant when compared to both the OLS and LRPS estimators.

Interestingly, when sVAR was tuned for optimal shrinkage, only relatively few off-diagonal edges were left in the estimated VAR matrix *Â*, i.e. not only was this matrix very sparse, but the structure the estimator highlighted as useful (in a predictive sense) was largely confined to the diagonal. In essence, one could achieve a similar performance to sVAR by simply adopting independent AR(1) models for each ROI, and ignoring ROI-interactions. Whilst diagonal structure was present systematically in the sVAR estimates, the region-region interactions (edges) were typically different across different subjects, and thus when averaging across subjects this resulted in very small values for *Ā*_*ij*_ when *i* ≠ *j*. This results in what appears to be very empty connectome plots in Figs. 6-7, however, the t-statistics do still indicate some off-diagonal structure, largely due to the standard-errors for these off-diagonal entries also being small. In contrast, the connectome plots for LRPS and VAR show much more widespread connectivity. In the limit where 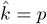 LRPS will recover the VAR model exactly, however, there is a large alignment in connectivity structure between LRPS and VAR even when only a few eigenvectors are used (*k* = 1, *k* = 10 in the examples here).

At a high level, the connectivity plots align somewhat with the activities that generated the datasets. Whilst individual differences make it hard to infer a ‘typical’ structure in the VAR matrix, we note that the overall level of connectivity, as evidenced by ave(*Â*), reduces in the resting-state dataset as compared with the task-based experiments. Furthermore, and likely associated with this reduction in connectivity, we note that the tuning parameters selected for LRPS automatically adapted to select a more parsimonious model in the case of the resting-state data, compared to the case of the task-based data. For example, we note in for the task-based datasets, our cross-validation procedure selects 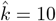, contrasting with the more restrictive 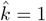 in the resting-state case. This could be a consequence of either the task-based data requiring a more expressive model, or the smaller sample-size in the resting-state dataset (*∼* 600 datapoints compared with *∼* 150) leading to overfitting if the model isn’t suitably restricted. Either way, it appears the cross-validation approach can be effective for gauging the level of adaption required. Notably, LRPS required quite a significant change in the tuning parameter when moving from task-based to resting-state data; however, sVAR used the same tuning parameter for both settings. The average connection in the task-based data is around 0.005 (for all methods), falling to 0.001 for the resting-state data.

Overall, we find that LRPS seems to increase the alignment between individual estimates and the population mean, as evidenced by smaller principal angles reported. This can potentially improve the interpretation of the population mean *Ā* in the LRPS case, however, we still note even when using LRPS there is significant variation, the angles are small compared to other methods, but not necessarily in absolute terms. Nevertheless, this reduction in principal angle does serves to highlight the constraints imposed by LRPS, and even though in this case each individual can select a different basis (set of eigenvectors) in which predictions are formed, the fact that we constrain the rank still encourages the estimated *Â*^(*l*)^ matrices to be better aligned geometrically, across the population.

## 5 Conclusion

In this article, we compare a low-rank based model with conventional methods as an alternative approach for modelling functional connectivity and Granger causality among ROIs. Compared to classical methods such as sVAR and VAR, LRPS offers a balanced trade-off between interpretability and predictive performance. The sVAR model excels at identifying subject-specific connectivity patterns by enforcing sparsity, effectively removing weaker connections, and performs well in brain activity prediction tasks. On the other hand, the standard VAR model, while less effective for prediction, tends to produce denser Granger causality patterns (when averaged over individuals) due to the absence of constraints. In contrast, our proposed LRPS method delivers competitive prediction performance with sVAR models whilst also allowing dense parameter matrices and more pervasive Granger causal relations, due to its low-rank formulation.

The performance of the proposed LRPS approach may be influenced by several factors. In particular, the choice of the rank parameter *k* plays a critical role, as an overly small value may oversmooth the data and remove meaningful temporal dynamics, while an overly large value may fail to adequately suppress noise. Additionally, departures from the assumed temporal dependence structure, such as nonstationarity or abrupt regime changes, may affect performance.

In this work, we deliberately did not pursue a joint low-rank and sparse modelling framework. Although joint low-rank and sparse models offer a flexible modelling paradigm, they were not considered here in order to maintain methodological simplicity and computational scalability. The joint models typically involve substantially higher computational complexity and additional tuning parameters, which can complicate both implementation and interpretation in high-dimensional settings. Exploring joint low-rank and sparse formulations within the proposed framework presents a natural direction for future research. In a scientific direction, it can be of interest to examine and compare Granger-causal structures in larger populations, for example, considering specific diseases and extending to larger datasets such as ADNI [62] to further validate the empirical behaviour of the estimation approaches, and how fMRI predictivity may be linked to pathology.

## Acknowledgements

We would like to acknowledge feedback provided by the Neuroscience at Lancaster (NeuRaL) research group on results presented in this work. Gemini and ChatGPT were used for some grammatical polishing on an early draft of this paper, however, these sections have largely been re-written by hand.

## Supporting information

**S1 Fig. Rank selection and prediction performance results for simulation experiment 1A and 1B**.

## Notes

### Competing Interest Statement

The authors have declared no competing interest.

